# Development of antigen-dextramers for detection and evaluation of CAR T cells

**DOI:** 10.1101/2024.11.27.625576

**Authors:** Rasmus U. W. Friis, Maria Ormhøj, Cecilie S. Krüger-Jensen, Markus Barden, Keerthana Ramanathan, Mikkel R. Hansen, Hinrich Abken, Sine R. Hadrup

**Author notes:** Contributed equally (Rasmus U. W. Friis, Maria Ormhøj).

## Abstract

**Background:** Chimeric antigen receptor (CAR) T cell therapy has transformed the treatment landscape of hematologic cancers by engineering T cells to specifically target and destroy cancer cells. Monitoring CAR T cell activity and function is essential for optimizing therapeutic outcomes, but existing tools for CAR detection are often limited in specificity and functional assessment capability.

**Methods:** We developed antigen-dextramers by conjugating multiple CAR-specific antigens to a dextran backbone. The dextramers were compared to previously reported antigen-tetramers for their ability to stain and detect CAR T cells. Because these multimers incorporate the CAR target antigen, they uniquely enable assessment of CAR T cell functionality by facilitating binding and activation analyses. We tested the staining and functional properties of the multimers across a range of CAR constructs with different affinities, using flow cytometry, microscopy, and NFAT-luciferase reporter assays.

**Results:** The antigen-dextramers demonstrated high specificity and sensitivity in staining CAR T cells, with adjustable antigen density to optimize binding. Antigen-dextramers also enabled effective clustering and subsequent activation of CARs, showing their utility as both a staining and functional assessment tool. The dextramers revealed that CARs with different affinities and clustering tendencies displayed varied binding and activation in response to different antigen densities.

**Conclusion:** Antigen-dextramers offer a dual advantage as versatile reagents for both staining and functional analysis of CAR T cells. Their capacity to engage CARs with the specific antigen provides a valuable platform for evaluating CAR functionality, informing CAR design improvements, and enhancing therapeutic precision.

## Background

Immunotherapy based on chimeric antigen receptor (CAR) T cells has proven highly successful against hematological cancers. CARs are synthetic receptors engineered to equip T cells with the ability to recognize and eliminate cancer cells. By combining an antigen-binding domain with intracellular signaling domains, CARs enable T cells to directly target cells expressing specific surface antigens and activate the T cell upon target engagement. This bypasses the need for traditional mechanisms of antigen presentation, offering a promising therapeutic option where conventional treatments have failed. The clinical success of CAR T cell therapy has encouraged scientists worldwide to extend it from hematological malignancies to solid tumors, autoimmune diseases, and infectious diseases. Consequently, several new CARs are under development, and many are currently being evaluated in clinical trials.

Often described as a “living drug,” in vivo expansion and persistence of CAR T cells are critical biomarkers linked to response, toxicity, and efficacy. Thus, monitoring CAR T cells post-infusion has become an integral part of most early-phase clinical trials. Both clinical and pre-clinical evaluations require reagents to stain CAR T cells for applications like multicolor flow cytometry, fluorescence-activated cell sorting, and microscopy. However, most of the existing CAR-staining reagents, such as polyclonal anti-IgG antibodies and Protein L, are limited by non-specific binding, incompatibility with other antibodies, and requirements for multi-step staining procedures. More specific CAR-staining reagents, like anti-idiotype antibodies, are often not commercially available and are limited by high developmental costs. Furthermore, most CAR-staining reagents are limited by ≤2-valency binding, and it has recently been demonstrated that higher-valency binding can improve CAR detection performance.

To address these limitations, we developed antigen-dextramers as high-avidity CAR-staining reagents. Dextramers combine multiple CAR target antigens on a dextran backbone, achieving multivalent binding that enhances CAR staining sensitivity. Beyond staining, however, dextramers uniquely enable the functional assessment of CAR T cells, as they mimic natural antigen engagement and facilitate CAR clustering on the cell surface. By adjusting the antigen density on dextramers, we can evaluate how different CAR designs interact with their target antigens, revealing functional insights related to CAR binding affinity, clustering behavior, and downstream activation. This dual utility makes dextramers valuable not only for CAR detection but also for assessing the functional properties of CAR T cells.

## Methods

### Cell lines and healthy controls

Peripheral blood mononuclear cells (PBMCs) from healthy donors were isolated from buffy coats provided by the Central Blood Bank of Copenhagen University Hospital (Rigshospitalet). The collection of donor material was conducted with the approval of the local Scientific Ethics Committee, and written informed consent was obtained from all participants in accordance with the Declaration of Helsinki. All cell lines were procured from the American Type Culture Collection (ATCC), USA.

### Lentiviral production and transduction

Four second-generation CARs were designed with either an anti-CD19 (FMC63)^11,13^, anti- CD79b (SN8)^14,15^, anti-CEA (BW431/26)^16^, or anti-HER2 (C6.5)^17^ scFv, a CD8 hinge and transmembrane domain, and intracellular domains from CD28, 4-1bb, and CD3ζ. Expression of the CAR was linked with expression of a green fluorescent protein (GFP) reporter gene. The CAR was inserted into a third-generation lentiviral vector under the control of a human EF1α promoter. All plasmids were synthesised by GenScript, USA.

Lentiviral particles were generated in HEK-293T cells by transfecting them with three separate packaging plasmids gifted by Didier Trono: pRSV-Rev (Addgene plasmid #12253; http://n2t.net/addgene:12253; RRID:Addgene_12253), pMDLg/p.RRE (Addgene plasmid #12251; http://n2t.net/addgene:12251; RRID:Addgene_12251), and pMD2.G (Addgene plasmid #12259; http://n2t.net/addgene:12259; RRID:Addgene_12259), along with a transfer plasmid containing the DNA construct of interest. The titer of the produced lentivirus particles was assessed through a titration experiment by infecting SUP-T1 cells and subsequently analysing GFP or mCherry expression via flow cytometry.

T cells were isolated from PBMCs of healthy donors using the EasySep™ Human T Cell Isolation Kit (STEMCELL Technologies). They were activated for 24 hours at 37°C and 5% CO2 in Hepes-buffered RPMI medium containing 20 IU/mL recombinant human IL-2 (Peprotech, USA) using anti-CD3/CD28-coated Dynabeads™ (Gibco™) at a 3:1 bead-to-cell ratio. After initial activation, the T cells were transduced with lentivirus at a multiplicity of infection (MOI) of 1 to 5 and expanded for another 10 to 14 days.

### Assembly of antigen multimers

Antigen dextramers were formed by combining allophycocyanin (APC) or phycoerythrin (PE)-labelled dextran backbone, conjugated to streptavidin molecules (FINA Biosolutions, USA), with mono-biotinylated protein antigens (Acro Biosystems, USA) at various molar ratios. In a similar manner, antigen tetramers were formed by gradually adding APC- or PE-conjugated streptavidin to mono-biotinylated antigen (adding 1/4 of the total amount of streptavidin every 5-10 minutes). Following a 20-minute incubation at 4°C, the multimers were supplemented with 10% freeze buffer, containing 0.5% BSA (Sigma-Aldrich, USA) and 5% glycerol (Fluka, USA) and stored at -20°C.

### Flow cytometry and cell sorting

Cells were centrifuged at 500 x g for 5 minutes at 4°C, and the supernatant was discarded. The pelleted cells were then resuspended in 5 µL of 1 µM Dasatinib (LC Laboratories, D- 3307) and incubated for 5-15 minutes at 4°C. After incubation, antigen multimers were added at desired concentrations, and the cells were incubated for 15 minutes at 37°C in the dark. The cells were washed once with FACS buffer, followed by staining with Near-IR (NiR) viability dye (Invitrogen™, L34976) and additional antibodies for surface staining for 30 minutes at 4°C in the dark. Finally, the cells were washed twice with FACS buffer, resuspended in 200-300 µL of FACS buffer, and immediately analysed on a LSRFortessa™ flow cytometer (BD, USA). Alternatively, the cells were fixed with 50 µL of 1% paraformaldehyde for 1-2 hours, washed twice with FACS buffer, and analysed 2-24 hours later.

Cell lines were sorted on a FACSAria™ flow cytometer (BD, USA) using a 100 µm nozzle. Amplitude was adjusted to optimise droplet break-off, and droplet calibration was performed before each sort using AccuDrop™ beads (BD, USA). CAR transduced cells were sorted based on GFP-expression, and sorting gates were set using untransduced cells. Cells were bulk sorted into tubes containing Hepes-buffered RPMI medium. After sorting, the cells were centrifuged at 500 x g for 5 minutes, resuspended in Hepes-buffered RPMI medium, and cultivated at 37°C with 5% CO2.

### Confocal microscopy

CAR-GFP SupT1 cells were stained with fluorescently labelled antigen multimers, and antibodies as described above. The cells were fixed using 1% Paraformaldehyde in PBS for 30 minutes at room temperature, and cell nuclei were stained for 10 min with 4′,6- diamidino-2-phenylindole (DAPI; Thermo Fisher, USA). Finally, cells were mounted on glass slides using ProLong™ mounting medium (Thermo Fisher, USA) and imaged using a LSM710 confocal laser microscope (Carl Zeiss, Germany). Image analysis was performed using Zeiss Zen software v3.1 (Carl Zeiss, Germany).

### Luciferase-based activation assay

CAR-mediated activation was assessed using anti-CD19 CAR-transduced reporter Jurkat cells (Signosis, USA), which produce luciferase in response to NFAT pathway activation. The transduced Jurkat cells were co-cultured with CD19-positive JEKO-1 cells for 15 hours. Following the co-culture period, the Jurkat cells were lysed using Pierce lysis buffer (Thermo Scientific, USA), and luciferin was subsequently added to the cell lysate. Luminescence was then measured using a microplate reader.

## Results

We have developed and evaluated fluorescently labelled dextramers and tetramers, conjugated to CAR antigens, for detection of CAR T cells. In short, CAR antigens were attached to fluorescently labelled dextran- or streptavidin molecules and used to stain human cell lines and primary T cells engineered to express CARs through lentiviral transduction (Figure 1 a). We validated the two multimers on four second-generation CARs: Anti-CD19, anti-CD79b, anti-CEA, and anti-HER2. The CAR constructs differed in their antigen-binding domains and signalling domains, while maintaining identical promoter sequences and overall structure. To confirm successful expression by flow cytometry, the CAR protein was linked to GFP (Figure 1 b).

**Figure 1.**
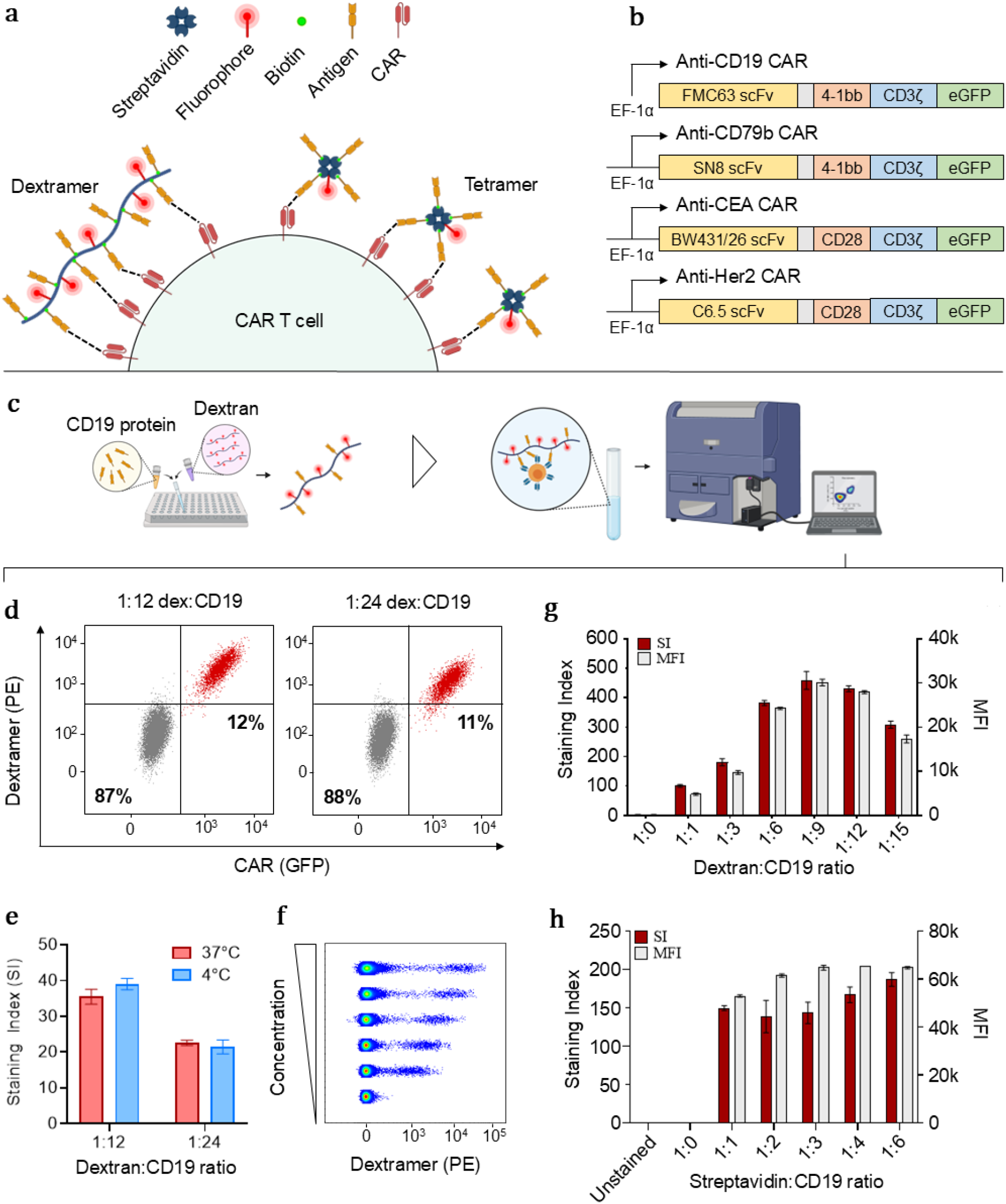
Development and optimisation of antigen-multimers using CAR-expressing cell lines. **(a)** Composition of antigen-dextramers and antigen-tetramers. **(b)** Configuration of CAR constructs used for validating antigen-multimers. **(c)** Overview of assembly process for generation of antigen-multimers. **(d)** Flow cytometry plots of ∼10% anti-CD19 CAR-expressing SupT1 cells stained with PE-labelled CD19- dextramers. **(e)** Influence of temperature during staining on the binding properties of CD19-dextramers (n=2). **(f)** Concentration-dependent fluorescent signal from anti-CD19 CAR-expressing SupT1 cells stained with increasing amounts of CD19-dextramer (1:9, dextran:CD19). **(g)** Titration of CD19 onto streptavidin-conjugated dextran backbone or **(e)** individual streptavidin molecules to assess the impact of different amounts of antigen on the staining of anti-CD19 CAR-expressing SupT1 cells (n=2).

### Optimisation of antigen-multimers using CAR-expressing cell lines

To get a thorough understanding of their staining properties, the antigen-multimers were initially tested on anti-CD19 CAR-transduced cell lines, including human Jurkat and SupT1 cells. CAR expression was identified by GFP luminescence. Transduced cell lines were sorted based on CAR expression and subsequently spiked into a population CAR-negative cells to obtain two well-defined cell populations. Biotinylated CD19 molecules were titrated onto PE-labelled dextran- or streptavidin molecules, resulting in multimers with an increasing number of antigens (Figure 1 c). The assembled multimers stained anti-CD19 CAR-expressing cells in a concentration-dependent and specific manner, at both 4°C and 37°C, while generating negligible non-specific fluorescence (Figure 1 d-f). To demonstrate the potential for extending these multimers to other CAR specificities, we developed CD79b-multimers to stain cells expressing anti-CD79b CARs, which exhibited similar staining properties (Supplementary Figure 1)^14,15^.

Based on the Mean Fluorescence Intensity (MFI) and the Staining Index (SI) of the multimer-binding populations, the number of antigens on the dextramers had a high impact on their binding capability. This was not as pronounced for the tetramers, likely due to their lower antigen valency and the high affinity of the CAR (Figure 1 g-h). In line with previous results reported by Hu et al., the high-valency dextramers exhibited superior staining, compared to the tetramers, indicated by a higher MFI and SI of the multimer-binding populations^11^. However, based on our results, increasing the number of antigens over 9 per dextran backbone resulted in loss of staining intensity, potentially due to steric hindrance.

### Antigen multimers exhibit high detection specificity and accuracy

The use of antigen multimers as CAR-staining reagents is highly dependent on their ability to selectively bind cells that express their cognate CAR. Therefore, we performed a cross- staining experiment to validate their binding specificity using three different CARs with varying affinity for their target: anti-CD19 (Kd = 0.3 nM)^11,13^, anti-CEA (Kd = 229 nM)^16^, and anti-Her2 (Kd = 16 nM)^17^. The results demonstrated that both antigen tetramers and dextramers stained ≥97% of their matched CAR-expressing cells, while staining of CAR- negative and non-matched CAR-expressing cells was negligible (<1%) (Figure 2 a-b).

**Figure 2.**
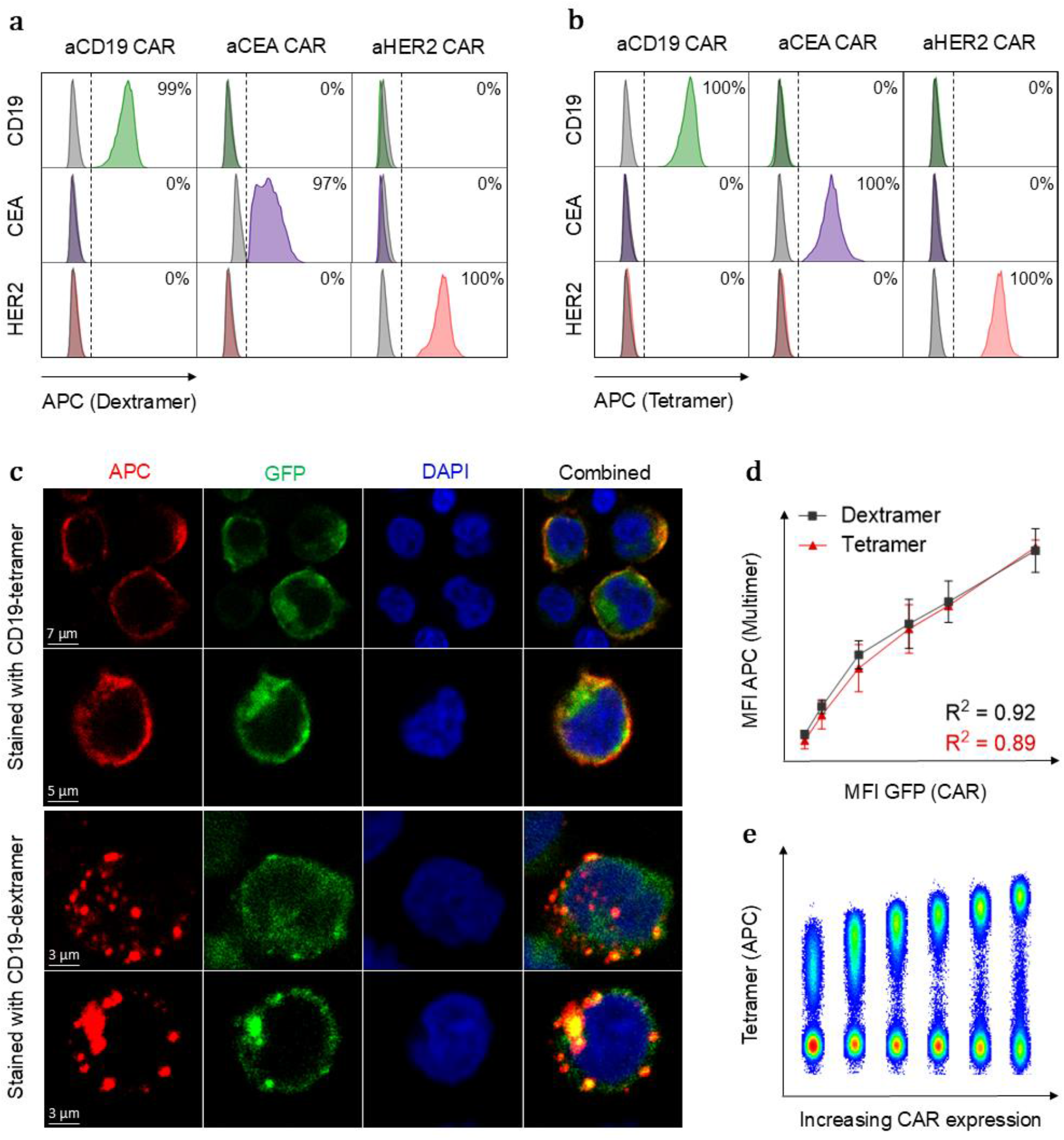
Dextramers and tetramers exhibit similar detection specificity and accuracy despite differences in their cell surface distribution patterns. **(a)** The ability of the antigen-dextramers and **(b)** antigen-tetramers to specifically detect CAR-expressing cells was assessed by cross-staining three Jurkat cell lines expressing different CARs. **(c)** Confocal microscopy images of anti-CD19 CAR-expressing Jurkat cells showing distinct cell surface distribution patterns of CD19-multimers. **(d)** Comparison between MFI of GFP (CAR) and APC (multimer) reflecting the ability of CD19-multimers to accurately detect CAR expression (n=2). **(e)** Flow cytometry plot of Jurkat cells expressing increasing levels of anti-CD19 CAR stained with CD19-tetramers.

Using confocal microscopy, we evaluated the distribution pattern of the two multimers on the surface of anti-CD19 CAR-transduced Jurkat cells. A distinct pattern was observed for each reagent; antigen-tetramers displayed a uniform distribution across the surface of the cells, while antigen-dextramers cluster together, forming aggregates on the cell surface. This clustering could be caused by a higher avidity interaction with the CARs, leading to localised binding hotspots (Figure 2 c).

We investigated how the staining intensity of the two multimers correlated with the CAR expression levels on the cell surface. For this purpose, we developed Jurkat cell lines with increasing levels of anti-CD19 CAR on surface. Using flow cytometry, we observed a positive correlation between the MFI of the antigen-multimers and the CAR expression levels of the cells, demonstrating that both types of multimers accurately reflect the cell surface CAR expression (Figure 2 d-e). To further validate the detection of low levels of CAR, we transduced primary human T cells with a low MOI of lentivirus and compared tetramer-positive cells to GFP-positive cells. As expected, the percentages of tetramer-positive and GFP-positive cells were similar, but the tetramer staining resulted in a more distinct separation between CAR-positive and CAR-negative cells, making it easier to differentiate between the two populations (Supplementary Figure 2).

### Antigen-independent clustering of high-affinity CARs inhibits multimer binding

A distinctive characteristic of antigen-dextramers is their modular nature, which allows adjustment of the number of antigens on the dextran backbone. We hypothesised that this capability could be beneficial for studying the relationship between antigen density and CAR affinity. To investigate this, we transduced SupT1 cells to express a panel of Her2 CARs with different affinities and assessed the staining properties of multimers with various antigen densities.

The anti-Her2 scFv panel was previously generated through site-directed mutagenesis and covers a 4-log affinity range, with Kd-values between 10^-7^ and 10^-11^ M, determined by surface plasmon resonance (Figure 3 a)^17^. All CARs shared the same modular configuration and CAR-transduced cells were sorted based on their CAR expression levels. Unexpectedly, we found an inverse correlation between CAR affinity and staining intensity for both Her2-dextramers and Her2-tetramers, as confirmed by flow cytometry and confocal microscopy, except that the lowest-affinity CAR stained less than the CAR with the second-lowest affinity (Figure 3 b-e).

**Figure 3.**
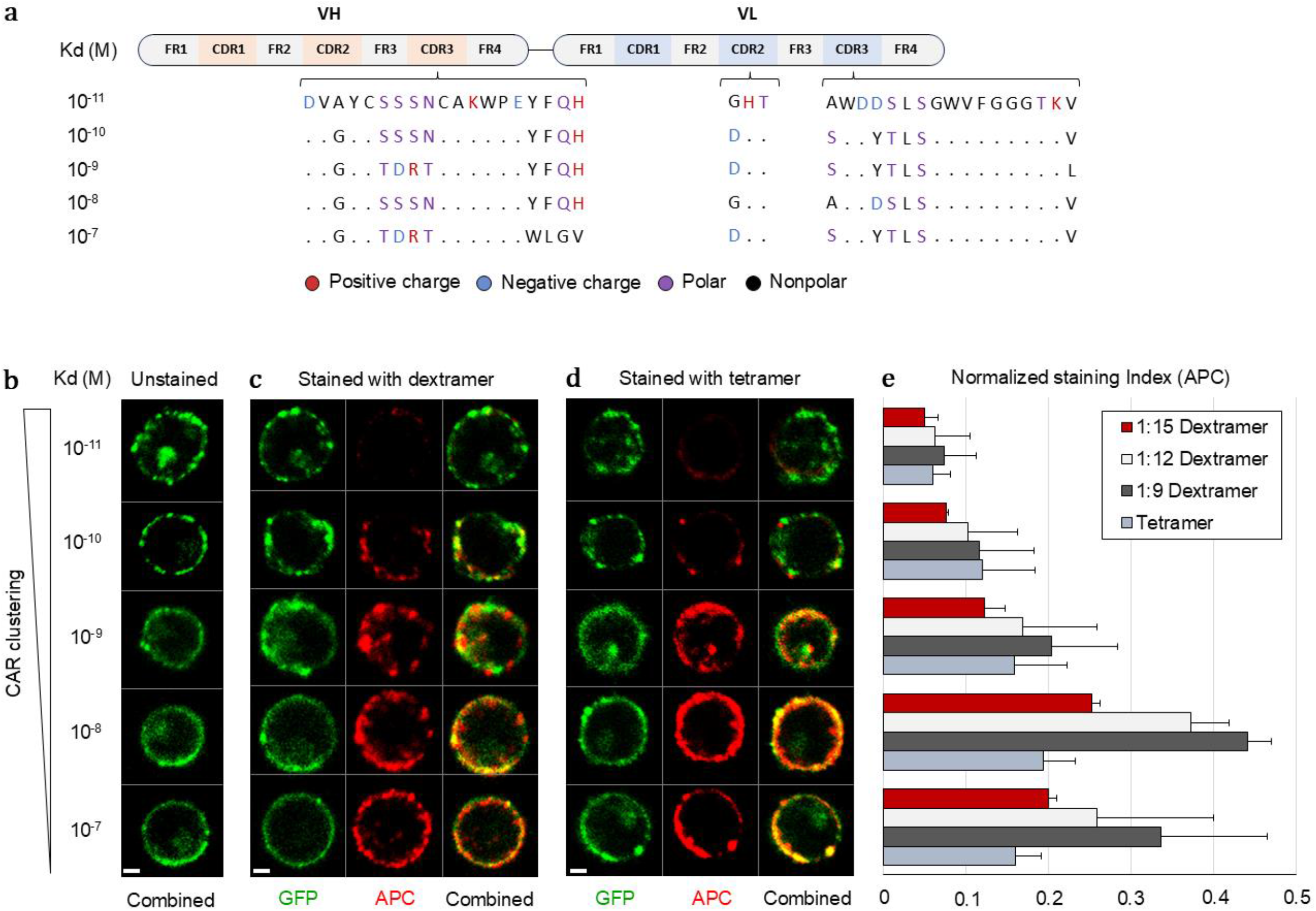
Antigen-independent clustering of high-affinity anti-Her2 CARs inhibits multimer binding. **(a)** Overview of anti-Her2 scFv panel showing amino acid substitutions in the CDR loops of the variable heavy (VH) and variable light (VL) chains. **(b)** Confocal microscopy images of SupT1 cells expressing anti-Her2 CARs with varying affinity stained with either **(c)** Her2-dextramers (1:9, dextran:Her2) or **(d)** Her2-tetramers revealing the impact of CAR clustering on multimer binding (scale bar=3 μm) **(e)** Comparison between the SI (normalised to CAR expression) of Her2-multimers with different amounts of antigen, used to stain SupT1 cells expressing the anti-Her2 CAR panel, determined by flow cytometry (n=2).

Further evaluation of the anti-Her2 CAR panel revealed that the high-affinity CARs tended to cluster spontaneously on the cell surface, even without antigen engagement (Figure 3 b). Based on this observation, we hypothesised that antigen-independent CAR clustering, due to scFv-intrinsic properties like homo- and oligomerisation, inhibits multimer binding.

### A tool for examining CAR-antigen interactions and their functional implications

We asked whether binding to tetramers or dextramers loaded with antigen result in downstream NFAT activation. We engineered NFAT-luciferase reporter Jurkat cells to express the panel of anti-Her2 CARs, exposed them to Her2-dextramers, and subsequently measured the luciferase activity in each cell culture. The results clearly demonstrated that CAR-transduced cells were activated in response to stimulation with antigen-dextramers, compared to unstimulated cells (Figure 4 a). Consistent with our previous observations, we found an inverse correlation between CAR affinity and CAR- induced activation, indicating that there is an optimum in CAR T cell activation by multimer binding, which decreases as CAR affinity increases, above a certain threshold where the affinity is too low. A similar pattern was observed when we stained with an anti- IgG antibody, although the antibody naturally did not reflect the difference in affinity (Figure 4 b). Additionally, we were able to co-stain the cells with antigen-multimers and anti-CAR antibody at the same time, which could be advantageous for specific applications (Supplementary Figure 3).

**Figure 4.**
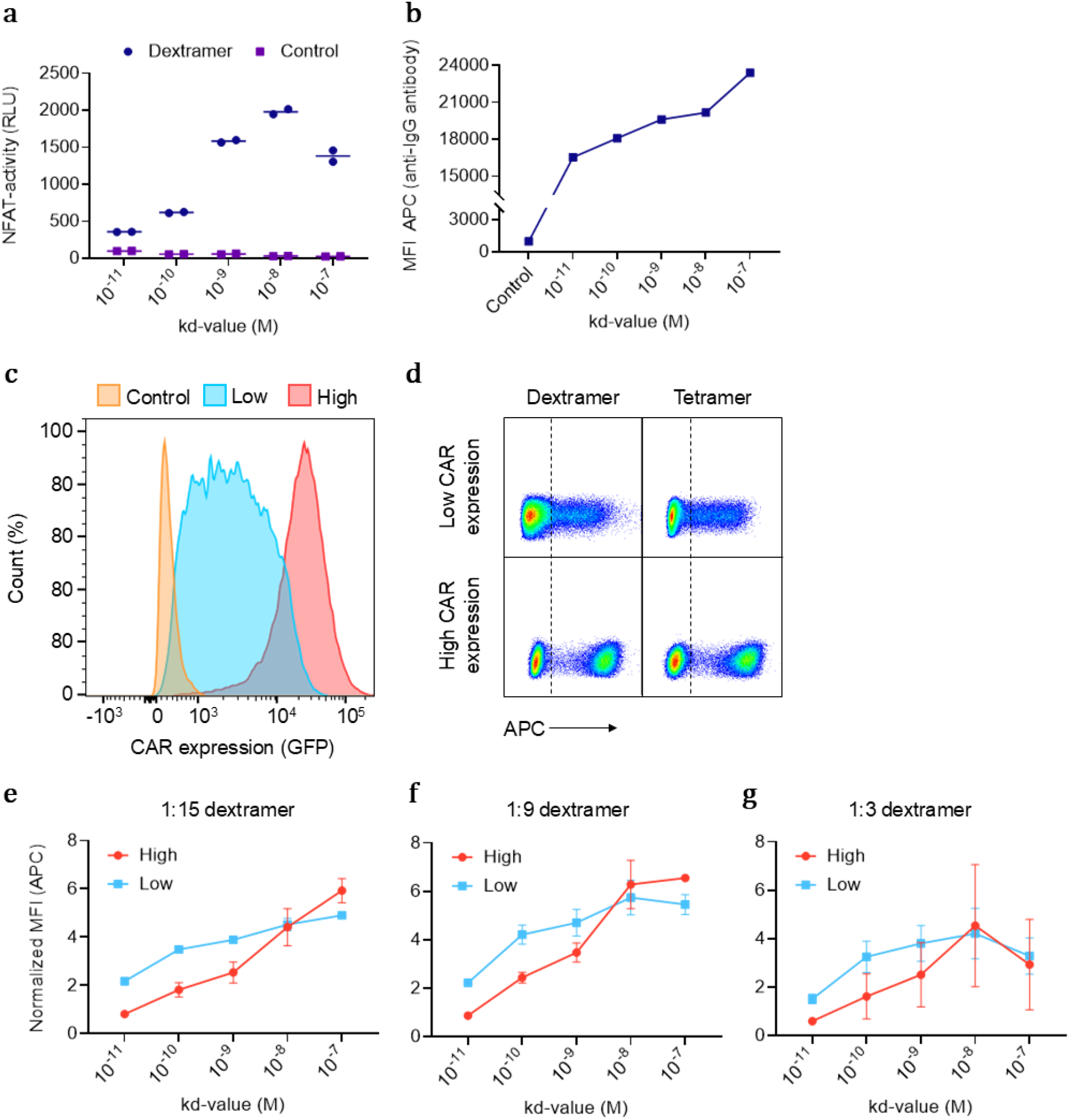
Antigen-multimers provide a tool for examining CAR-antigen interactions and their functional implications. **(a)** CAR-mediated activation of anti-Her2 CAR-expressing Jurkat cells after incubation with Her2-dextramers for 15 hours, measured based on NFAT-induced luciferase activity (RLU, relative light units). **(b)** Detection of anti-Her2 CAR-expressing Jurkat cells using an APC-labelled anti-IgG antibody (MFI, mean fluorescence intensity). **(c)** Anti-Her2 CAR expression levels in Jurkat cells after sorting into low- and high CAR-expressing populations. **(d)** Flow cytometry plot of Jurkat cells expressing low or high levels of anti-Her2 CAR after staining with Her2-multimers. **(e)** Impact of anti-Her2 CAR expression level on the binding properties of dextramers with a high dextran:Her2 ratio (1:15), **(f)** medium dextran:Her2 ratio (1:9) and **(g)** low dextran:Her2 ratio (1:3) (n=2).

Next, we investigated how reduced CAR expression levels influenced the binding properties of the Her2-dextramers. Jurkat cells were sorted into high and low CAR-expressing populations and stained with dextramers containing different amounts of Her2 (Figure 4 c-d). By calculating the SI normalised to the CAR expression for each cell line, we compared the relative dextramer binding between the high and low CAR-expressing populations (Figure 4 e-g).

Interestingly, the lower-affinity CARs (except for the lowest-affinity) exhibited better binding to dextramers in both high- and low-sorted CAR-expressing populations, with particularly enhanced binding at high expression levels. In contrast, the higher-affinity CARs showed better binding at low CAR expression levels, likely due to reduced spontaneous clustering that facilitates better dextramer access. Among the three dextramers, the one with the highest number of Her2 molecules (1:15) demonstrated the best binding to the low-affinity CARs compared to dextramers with fewer Her2 molecules (1:9 and 1:3). This indicates that higher antigen density can compensate for lower CAR affinity by maintaining sufficient antigen interaction despite weaker individual binding strength. Conversely, the dextramer with the lowest number of Her2 molecules showed the poorest staining for the lowest affinity CARs across both high and low CAR expression populations.

## Discussion

In this study, we developed and evaluated antigen-tetramers and antigen-dextramers; two high-avidity CAR-staining reagents based on biotinylated CAR antigens conjugated to fluorescently labelled streptavidin molecules. Through staining titrations and spike-in experiments, we demonstrated that these antigen-multimers detect CARs with high specificity and sensitivity in both CAR-transduced cell lines and primary human T cells. Antigen-tetramers contain four CAR antigens on a single streptavidin molecule, while antigen-dextramers consist of a dextran backbone with multiple streptavidin molecules, making them highly modular and extensible. This unique capability makes antigen- dextramers a valuable tool for evaluating new CAR designs based on their antigen binding properties. By systematically varying antigen density, researchers can assess how different CAR constructs interact with their corresponding antigens immobilized on a matrix. As exemplified in this study, this approach allowed for the comparative analysis of antigen density and CAR affinity, providing insights into the effects of CAR clustering behaviour, which can be used for rational optimisation of new CAR designs.

While it is generally assumed that higher affinity would lead to stronger responses, research on T cell activation suggests a more complex relationship. Both very high and very low affinities can lead to attenuated responses, indicating a non-linear (bell-shaped) relationship between affinity and T cell function^21,22^. In line with this, anti-CD19 CAR T cells with an intermediate affinity scFv (Kd of 14 nM) demonstrated higher proliferation and anti-tumour activity *in vivo* compared to those with a high affinity CAR (Kd of 0.3 nM)^23^. Although the effects of affinity and antigen density are well-documented for TCRs, these factors remain largely understudied in the context of CAR T cells.

Studies on pMHC-tetramers revealed that their binding affinity threshold was higher than needed for T cell activation, which means they potentially miss antigen-specific T cells with low-affinity TCRs that are common in both tumour-specific and autoimmune T cells^24^. Since CARs typically have high affinity for their target antigens, this might not translate to detection of CAR T cells. However, recent strategies suggest reducing CAR affinity to just above the activation threshold to minimise side effects while retaining anti- tumour efficacy^23,25^. These reduced-affinity CARs, currently in early clinical trials, could limit the detection sensitivity of antigen-multimers, making other detection methods more suitable. However, such "affinity-tuning" is challenging and empirical, as changes in affinity do not correlate linearly with activation strength^26^. Therefore, we anticipate that antigen-dextramers, with their ability to easily adjust the antigen density, could become a valuable tool for detecting and evaluating new low-affinity CARs during development.

Previous studies done by Dolton et al. showed that pMHC-dextramers stained antigen-specific T cells much brighter than the corresponding tetramers^27^. In line with this, Hu et al. demonstrated that high avidity staining reagents outperform mono- or bivalent staining reagents for detection of CAR T cells^11^. Consistent with these findings, we demonstrated that antigen-dextramers exhibited better staining properties compared to antigen-tetramers, indicated by a higher MFI and SI. However, the two multimers were equally capable of capturing almost all CAR-expressing cells with minimal staining of CAR-negative cells.

We successfully applied the antigen multimers on four different second-generation CARs: anti-CD19, anti-CD79b, anti-CEA, and anti-HER2 CARs. Our results suggest that the increased binding avidity of the antigen-dextramers leads to clustering on the cell surface, which amplifies the fluorescence signal and thereby enhances detection sensitivity. In theory, this clustering effect may limit detailed mapping of cell surface CAR expression, suggesting a trade-off between signal amplification and spatial resolution. Therefore, antigen-tetramers, with their uniform distribution, could be more suitable for applications requiring detailed and comprehensive mapping of CAR expression, while antigen-dextramers, due to their clustering, might be better suited for applications needing enhanced signal intensity.

Additionally, we demonstrated that antigen-dextramers can activate CAR T cells *in vitro*. Unlike generic T cell mitogens, antigen-dextramers are CAR-specific and our results suggest that they mimic CAR stimulation through CAR oligomerisation and the formation of an artificial immunological synapse^32,34,35^. We hypothesise that specialised antigen- dextramers containing co-stimulatory molecules and cytokines, similar to the recently reported pMHC scaffolds, can be used for *ex vivo* expansion of CAR T cells. This offers an alternative to current methods that rely on anti-CD3/CD28 beads or irradiated antigen- positive feeder cells^36,37^.

Antigen-multimers offer distinct advantages for detecting CAR T cells by microscopy and flow cytometry. Unlike anti-IgG antibodies and Protein L, antigen-multimers can stain CAR T cells alongside exogenous antibodies, making them highly suitable for multicolour flow cytometry. Additionally, antigen-multimers are not limited to specific anti-idiotypic antibody clones but can bind any CAR targeting a given antigen. For example, anti-FMC63 anti-idiotype antibodies can only stain FMC63-based CARs, which excludes other anti- CD19 CARs like fully human anti-CD19 CARs or new CARs engineered with lowered binding affinity^23,38,39^. Because they are not restricted to specific antibody clones, antigen- multimers can easily be adapted for use with both existing and new CAR systems, or even new synthetic receptor designs like the HLA-independent T cell (HIT) receptor, which has recently been proposed to overcome tumour escape associated with low target antigen expression^40^. Finally, bi- and tri-specific CARs are being evaluated in preclinical and early clinical trials to address the issues of tumour heterogeneity and antigen escape, and trials with two CAR T cell products have recently been initiated^41,42^. While most CAR staining reagents are unable to differentiate between different CAR constructs, antigen- multimers provide the capability to evaluate how each CAR interacts with its target antigen.

While the antigen-multimers demonstrated good specificity and sensitivity, the potential for steric hindrance at high antigen densities on dextramers suggests that further optimisation may be needed to maximise their binding efficiency across different CAR constructs and CAR expression levels. Additionally, investigating the long-term stability and reproducibility of these reagents will be essential for their widespread adoption in both research and clinical practice.

## Declarations

## Contributors

RUWF, MO, and SRH designed the research and wrote the manuscript. RUWF, MO, CSKJ, MB, KR, MRH, and SRH contributed to experiments and analysis. MB and HA provided important materials, methods and advice. SRH is responsible for the overall content as the guarantor.

## Data availability statement

All data relevant to the study are included in the article or uploaded as supplementary information. All data is present in this research article and Supplementary Materials, which includes Supplementary Figure S1–S5.

