## Supplementary Figures for "Development of antigen-dextramers for detection and evaluation of CAR T cells"

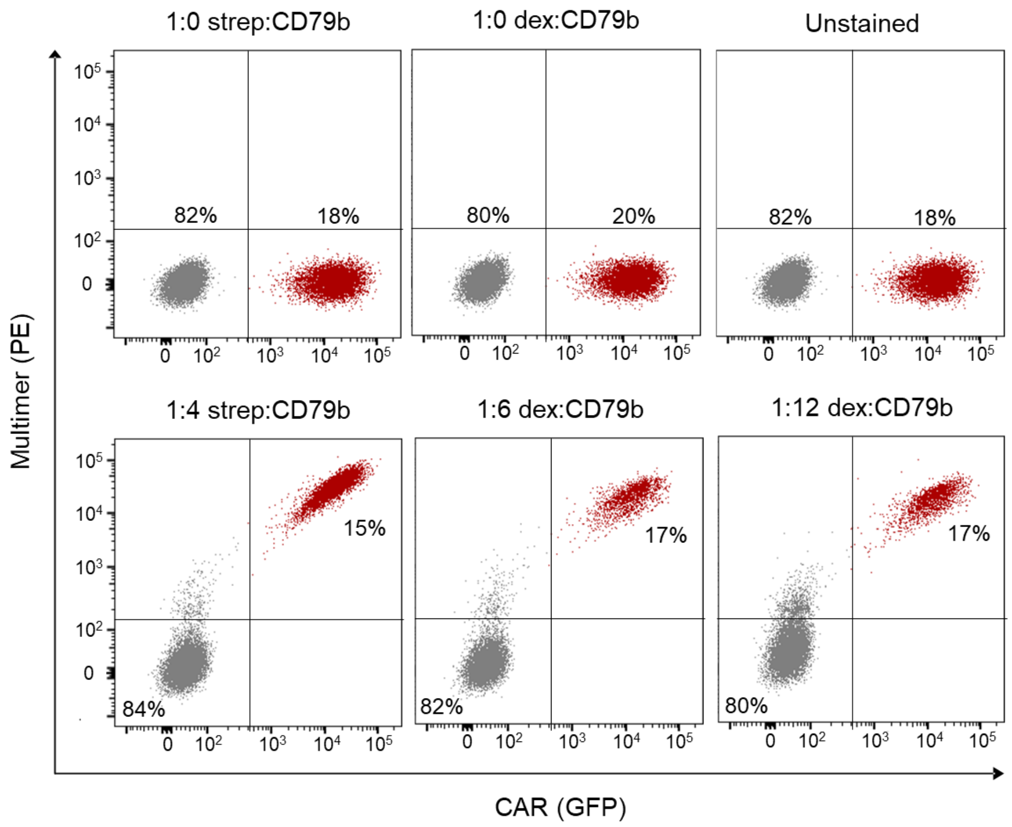


**Supplementary Figure 1. Antigen-multimer staining of anti-CD79b CAR.** Anti-CD79b CAR-expressing SupT1 cells stained with PE-labelled CD79b-tetramer and two different CD79b-dextramers. Unstained cells and PE-labelled streptavidin or dextran backbones were used as controls.

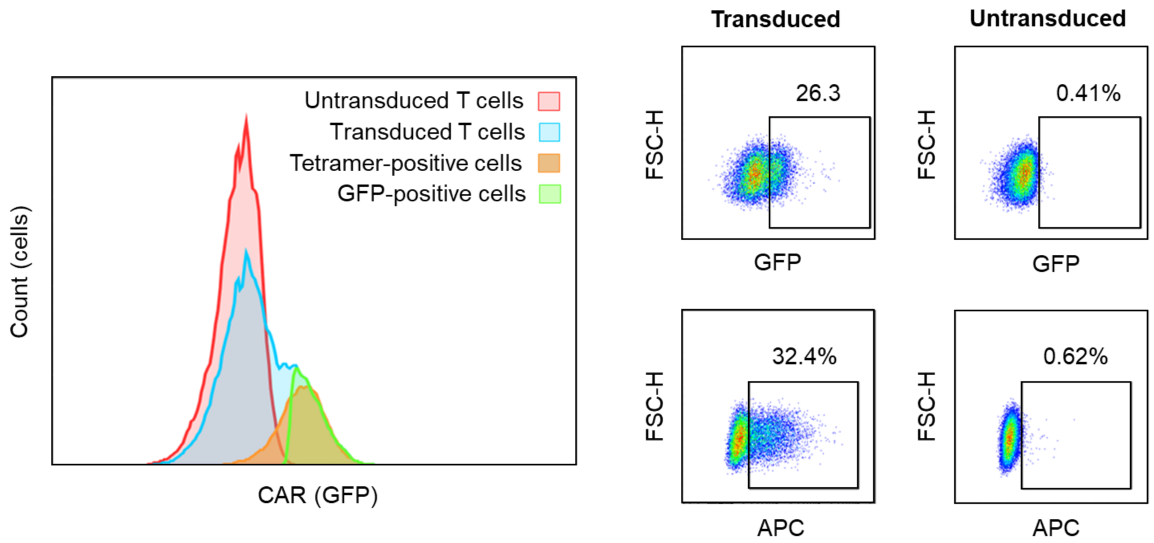


**Supplementary Figure 2. Antigen-tetramers detect low levels of CAR expression in T cells.** Comparison between GFP- and tetramer-based detection of anti-CD19 CAR-expressing T cells transduced with a low MOI of lentivirus. Gating was done manually using the FlowJo software.

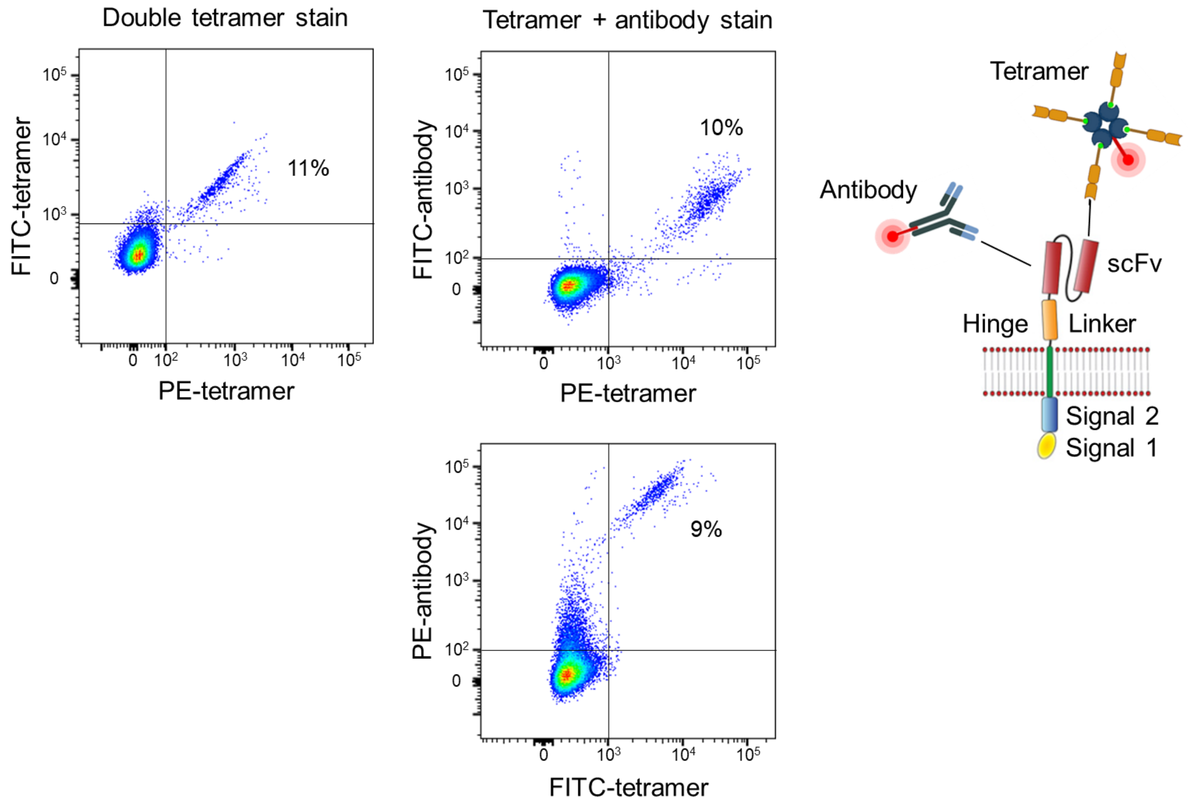


**Supplementary Figure 3. Antigen-tetramers can co-stain CARs with anti-IgG antibodies.** Staining of anti-Her2 CAR-expressing Jurkat cells demonstrating simultaneous binding of Her2-tetramers and anti-IgG antibodies.
